# Selective degradation of oncogenic extrachromosomal DNA by type I-E CRISPR

**DOI:** 10.1101/2025.09.21.677573

**Authors:** Ensieh M. Poursani, Vu v. Pham, Anahid Ehteda, Orazio Vittorio

## Abstract

Extrachromosomal DNA (ecDNA) is a major driver of oncogene amplification, intratumoral heterogeneity and therapy resistance across multiple cancer types, yet there are currently no effective strategies to selectively eliminate it. Here, we show that the type I-E CRISPR system can specifically target and degrade ecDNA in human cancer cell lines (COLO320DM and GBM39). By designing guide RNAs targeting ecDNA-specific breakpoints absent from chromosomal DNA, we achieved efficient depletion of MYC- and EGFR-containing ecDNAs in COLO320DM and GBM39 cells. Loss of ecDNA was accompanied by diminished oncogene signaling, disrupted ecDNA architecture, and impaired tumor cell proliferation, without detectable chromosomal off-target activity. These findings establish a proof-of-concept framework for directly targeting oncogenic ecDNAs and highlight type I-E CRISPR as a promising platform for therapeutic development in ecDNA-driven cancers.

## Main

ecDNA is an overlooked but critical driver of tumor heterogeneity and therapeutic resistance^1-3^. Unlike chromosomal DNA, ecDNA exists as circular, acentromeric fragments that replicate independently and segregate unevenly during cell division^4-7^. This non-Mendelian inheritance contributes to high oncogene copy number variation and promotes intratumoral genetic diversity^4,8,9^. ecDNAs often harbor amplified oncogenes and regulatory elements, enhancing their expression and accelerating tumor progression^10-14^. Moreover, ecDNAs exhibit high mutation rates, dynamic structural rearrangements, and can cluster into transcriptionally active hubs—collectively driving rapid genome remodeling^11,15^. These features enable cancer cells to adapt swiftly to selective pressures, making ecDNA a central engine of genome plasticity and a major contributor to treatment resistance. Despite its significance, ecDNA remains a largely untargeted vulnerability in current cancer therapies^16^. CRISPR technologies have transformed genome engineering, yet their application to ecDNA has been limited. While Cas9 and Cas12 nucleases create targeted double-strand breaks, the type I-E CRISPR–Cas3 system offers a unique capacity for long-range, processive DNA degradation^17^. Originally described as a programmable DNA interference (DNAi) device in bacteria^18^, type I-E CRISPR has recently been adapted for mammalian cells, where it induces unidirectional deletions spanning kilobases with minimal off-target activity^19^. Building on this property, we hypothesized that type I-E CRISPR could be harnessed to selectively target and degrade oncogene-bearing ecDNA in cancer cells.

The type I-E CRISPR system functions as a multi-protein complex that uses a CRISPR RNA-guided surveillance mechanism to recognize specific DNA sequences and then recruits the Cas3 nuclease-helicase^20^. Cas3 drives extensive, processive, and unidirectional degradation of DNA, effectively “chewing up” long stretches beyond the initial cleavage site^19^.

While CRISPR methods like Cas9 are widely used for precise genome editing, their activity is typically limited to making localized DNA cuts. In contrast, the type I-E CRISPR system enables broad and continuous DNA degradation via Cas3, allowing for comprehensive removal of target sequences such as oncogenic ecDNA with high specificity and minimal off-target effects^19^. This unique capability makes type I-E CRISPR a promising tool for applications demanding large-scale DNA elimination^19,21^.

To evaluate the effectiveness of this system, we employed type I-E CRISPR to specifically target and degrade plasmids, serving as a representative model of circular ecDNA. The plasmids pCMV6-AC-GFP and pmCherry-C1 were used as circular DNA targets for the CRISPR system. Guide RNAs were designed to target the Neomycin resistance gene, located approximately 1.5–2.5 kb away from the reporter genes GFP and mCherry, respectively. HeLa cells were co-transfected with the type I-E CRISPR components, guide RNAs, and the reporter plasmids. After 48 hours, cells receiving guide RNAs showed a significant reduction in fluorescence compared to mock controls (Fig. 1B & 1D). Next, COLO320DM and GBM39 cell line used as models of ecDNA-positive (ecDNA□) cancer cells. Guide RNAs targeting specific ecDNA breakpoints in COLO320DM and GBM39 were designed and cloned into an empty U6 vector. These ecDNA□ cells were then transfected with the type I-E CRISPR system along with pooled guide RNAs. Four days post-transfection, cells were stained with c-MYC (COLO320DM), and EGFR (GBM39) DNA FISH probes and fluorescent signals were captured using microscope.

**Figure 1.**
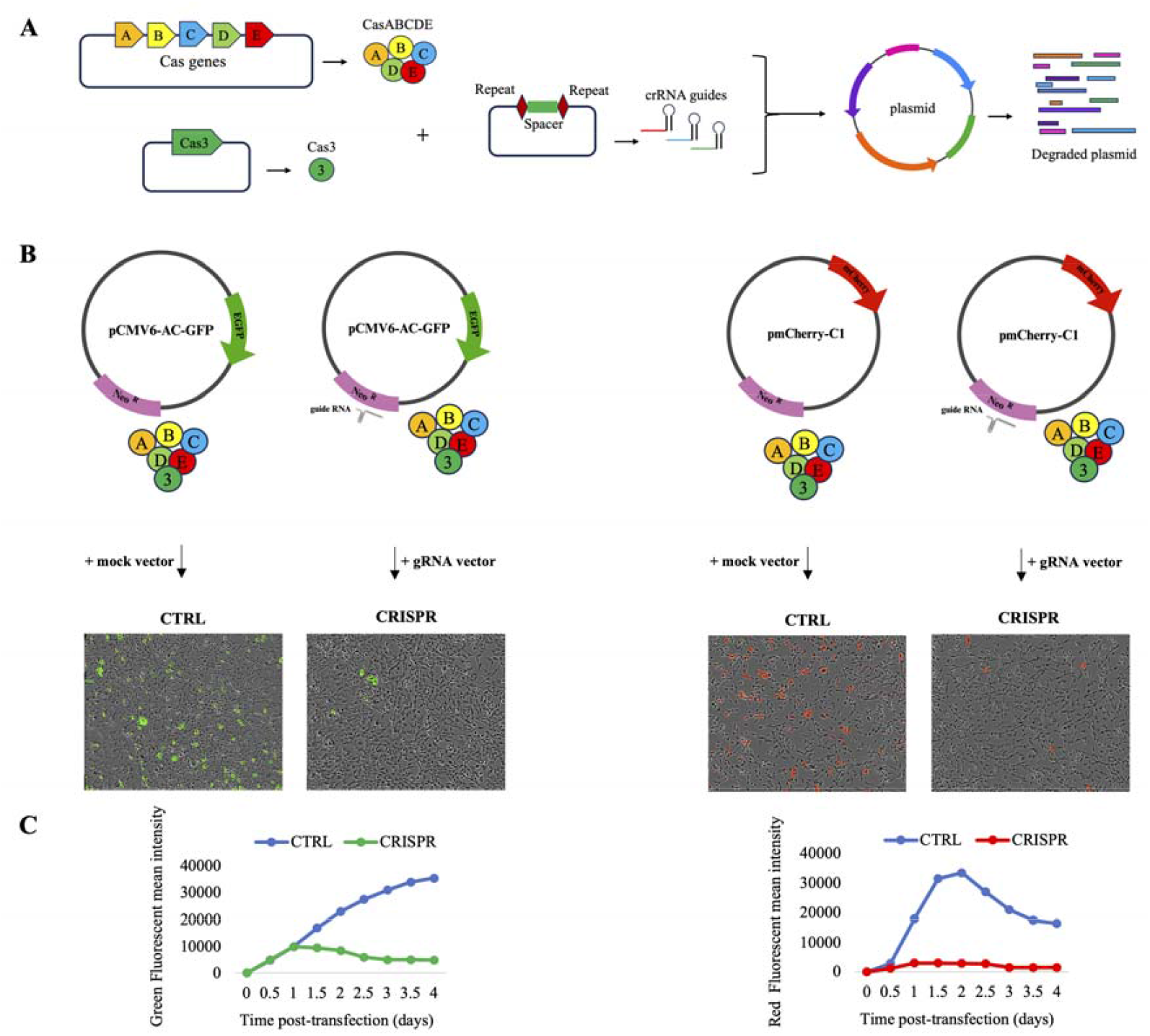
Workflow of type I-E CRISPR degrading circular DNA models (plasmids). **A)** A schematic representation of the type I-E CRISPR system. **B)** Targeting and degradation of DNA plasmids by type I-E CRISPR in Hela cells. Cells and fluorescent signals visualized by IncuCyte SX5 microscope. **C)** Time-course graphs illustrate the progressive degradation of target plasmids, as evidenced by a steady reduction in fluorescent signals. CTRL; mock control, CRISPR; guide RNAs against Neomycin resistance gene (Neo gRNA).

Our analysis revealed a clear reduction in fluorescent signals in cells treated with guide RNAs compared to mock controls (Fig. 2). Furthermore, comparison between treated and control cells showed structural alterations in ecDNA, likely resulting from CRISPR-mediated cleavage and/or degradation (Suppl. Fig. 2). Further studies are required to clarify the underlying mechanisms. Moreover, no off-target chromosomal effects were detected in metaphase spreads of either COLO320DM (ecDNA□) or MCF-7 (ecDNA□) cells (Suppl. Fig. 5). Finally, ecDNA degradation significantly reduced cell proliferation. Targeting ecDNA with specific guide RNAs led to a significant reduction in cell growth compared to mock-treated controls, highlighting the potential of targeting oncogenic ecDNAs to suppress tumor cell growth (Suppl. Fig. 3). Extrachromosomal DNA (ecDNA) plays a pivotal role in tumor progression and is linked to poor prognosis across multiple cancer types. These circular, acentric DNA elements frequently carry amplified oncogenes, driving elevated gene expression and contributing to aggressive tumor behavior. Despite the well-established role of ecDNA in cancer evolution and treatment resistance, no therapies currently exist that specifically target ecDNA. In this study, we demonstrate that the type I-E CRISPR system can selectively degrade ecDNA in cancer cells by targeting ecDNA-specific sequences. This approach effectively reduced oncogene copy numbers while preserving overall cell viability. Notably, ecDNA degradation resulted in decreased proliferation of ecDNA-positive cancer cells, highlighting its potential therapeutic value. By directly eliminating oncogene-carrying ecDNA, this strategy may provide a novel alternative to conventional treatments such as chemotherapy and radiotherapy, which are often associated with significant toxicity.

**Figure 2.**
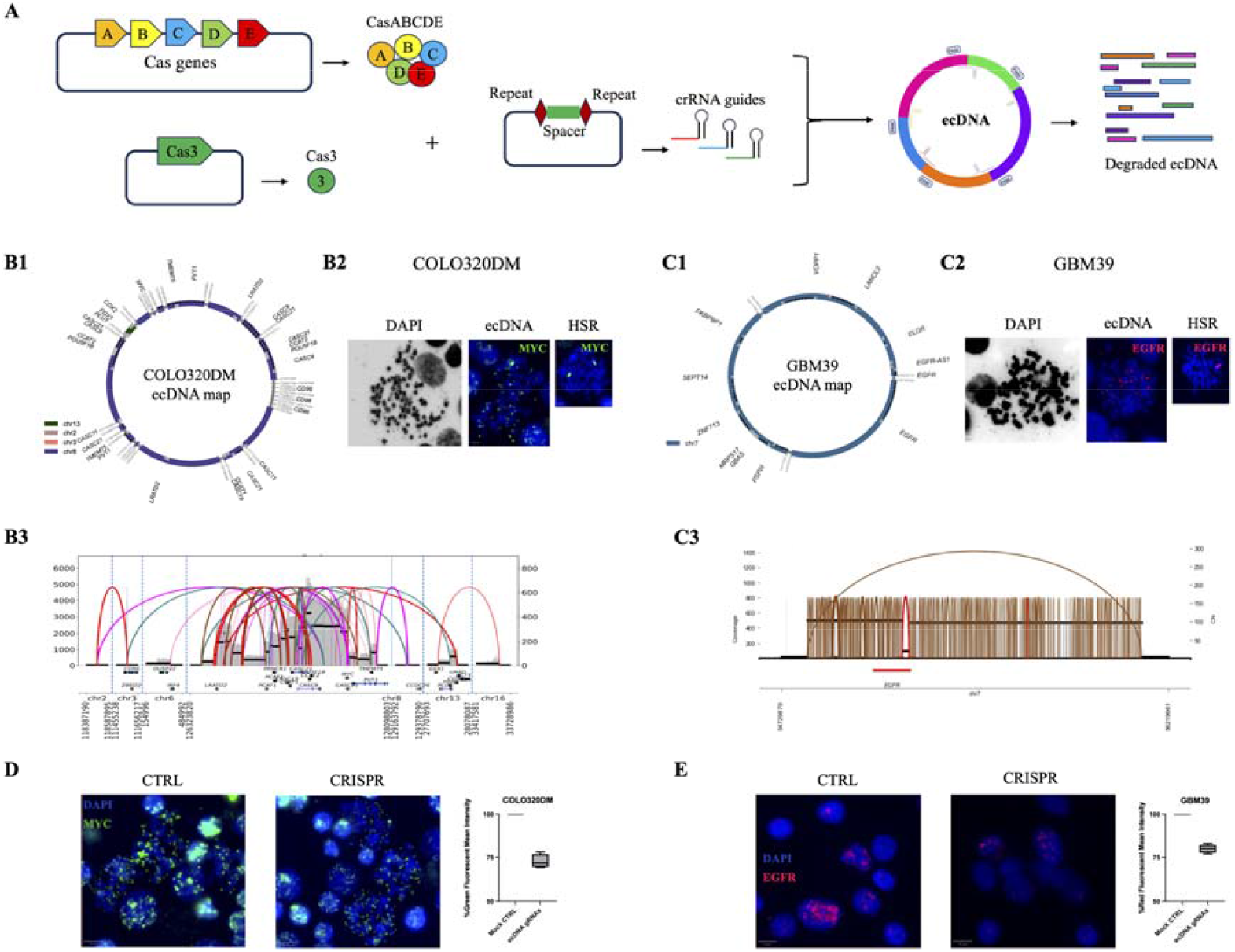
Workflow of type I-E CRISPR degrading ecDNA. **A)** A schematic representation of the type I-E CRISPR system selectively target and eliminate ecDNA. **B1 & C1)** EcDNA maps generated by CoRAL (COLO320DM) and AmpliconArchitect (GBM39); **B2 & C2)** metaphase spreads of COLO320DM and GBM39. Scale bas, 5µm; and **B3 & C3)** a structural variant (SV) view depicting the AmpliconArchitect (AA) reconstruction of the MYC amplicon in COLO320DM and EGFR amplicon in GBM39 cells. **D & E)** Representative MYC FISH in COLO320DM and EGFR FISH in GBM39 cells imaged by Confocal Microscope. Scale bar, 10µm. All imaging experiments were repeated at least three times, with similar results. CTRL; mock control, CRISPR; guide RNAs against ecDNA breakpoints.

Conceptually, this system could be extended to treat other diseases, including viral infections. We propose that the type I-E CRISPR system may be effective in targeting and degrading DNA viruses such as HCMV, HSV, EBV, and HPV. However, additional studies are necessary to confirm its therapeutic potential in these settings. While CRISPR-based degradation of ecDNA holds significant promise, its clinical translation faces key challenges—including efficient, tumor-specific delivery, potential immune responses, and the risk of off-target effects. Addressing these limitations will be essential to move CRISPR-based ecDNA targeting closer to clinical application.

In brief, our study establishes type I-E CRISPR as a novel approach for eliminating non-chromosomal DNAs including ecDNA. Ultimately, the ability to selectively eliminate ecDNA opens a new therapeutic avenue for cancers driven by oncogene amplification, with broad implications for both tumor biology and clinical intervention.

## Material & Methods

### Cell culture and treatments

Human colon cancer cell line COLO320DM and glioblastoma cell line was kindly provided by Prof. John Mariadason from Olivia-Newton John Cancer Research Institute (Australia) and cultured in DMEM with 10% fetal bovine serum (FBS). Human glioblastoma GBM39 tumour spheroid was kindly provided by Dr. Sameer Greenall from Monash University (Australia), and cultured in RHB-A cell culture media supplemented with with 1% GlutaMAX, B27, EGF, FGF and heparin. All cell lines tested negative for mycoplasma.

For the transient transfection procedure, we began by seeding 150,000 cells into each well of a 24-well plate. The cells were then transfected with a mixture containing 250 ng of Cas3, 250 ng of CasABCDE, 250 ng of NucS/ExoIII, and 1 µg of mixed guide RNAs for the treated group. In contrast, the mock control group received the same amounts of Cas3, CasABCDE, and NucS/ExoIII but was supplemented with 1 µg of an empty vector for guide RNA. Lipofectamine 3000 was utilized for the transient transfection process according to the manufacturer’s instructions.

### CRISPR Plasmids

The Cas3 and Cascade components, specifically Cas5, Cas6, Cas7, Cas8, and Cas11 from E. coli K-12, were acquired from Addgene with a bipartite nuclear localization signal (bpNLS) at both the 5′ and 3′ ends (Addgene ID: 134920 & 134919). The crRNA expression plasmid was constructed using crRNA sequences that included two BbsI restriction sites within the spacer region under the U6 promoter, synthesized by Integrated DNA Technologies (IDT). All targeted sequences are detailed in Supplementary Table 1. The crRNA expression plasmids were generated by inserting 32-bp double-stranded oligonucleotides corresponding to the target sequences at the BbsI restriction site (Addgene ID: 134921).

### DNA FISH

Coverslips containing fixed cells in metaphase were equilibrated with 2× SSC buffer, followed by dehydration in ascending ethanol series (70%, 85%, 100%) for 1 min each. DNA FISH probes (Empire Genomics) were added onto a slide, and the coverslip was applied. The FISH probe and sample were co-denatured on a 78□°C hotplate for 2 min, and hybridization was carried out overnight at 37□°C in a humidified chamber. The coverslips were removed and washed in 2× SSC, and then mounting medium including DAPI (VECTASHIELD) was applied and the coverslip was mounted onto a glass slide.

### Microscopy

Confocal Microscope (Leica Stellaris 8) and Olympus v200 were used to capture images using 63× oil lens rom each visual field, and images were analysed using ImageJ and Qupath at default settings (EGFR (red), MYC FISH (green), and DAPI (blue)).

### Oxford Nanopore Sequencing

Whole genomic DNA was isolated from COLO320DM cells using QIAGEN DNeasy Blood & Tissue kit, based on the manufacturer’s instruction. initial sample QC was done using Qubit and Genomic DNA screen tape on Tape station. to check the quality.

Ligation sequencing (SQK-LSK114) library prep kit was used for Library preparation. 12µL of final library (∼230ng) was loaded on PromethIon R10 flowcell used for sequencing and ran at high accuracy base calling for 72 hours.

### Identify extrachromosomal DNA (ecDNA) genome structures

We used coRAL (version 1.0.r1) (Zhu et al., 2024) to reconstruct ecDNA architectures using our long-read whole-genome sequencing data. The samples were sequenced using the Oxford Nanopore (ONT) ligation sequencing kit V14 (SQK-LSK114). The sequencing data was basecalled with high-accuracy basecalling (model v4.3.0, 400bps) using Dorado (version 7.4.12). The basecalled data was then aligned to the human reference genome (hg38) based on the wf-alignment workflow (https://github.com/epi2me-labs/wf-alignment). We utilised CNVkit (version 0.9.12) (Talevich et al., 2016) to compute the genome-wide copy number (CN) profile, which was used by CoRAL to identify and filter CN gain regions where amplifications existed. CoRAL was then applied to perform a breakpoint graph construction and cycle decomposition on the CN gain regions to identify candidate ecDNA structures. The outputs of CoRAL were visualised in Circos-style images with CycleViz (version 0.2.0) (https://github.com/AmpliconSuite/CycleViz)^22,23^.

## Acknowledgements

This research was funded by Hearts and Minds Foundation & TDM. We acknowledge Dr Sameer Greenall (Monash University) for providing the GBM39 cell line, Professor John Mariadason (Olivia Newton-John Cancer Research Institute) for the COLO320DM cell line, and Dr Prabh Phalora (University of New South Wales) for supplying DH5α competent cells.

## Author information

### Authors and Affiliations

### Contributions

E.M.P contributed to the study design and collection and interpretation of the data and wrote the manuscript. V.V.P conducted bioinformatic analysis. A.E. contributed by assisting with tissue culture and providing essential laboratory materials. O.V. supervised the project. All authors have read and agreed to the published version of the manuscript.

### Corresponding author

Correspondence to Orazio Vittorio

## Ethics declarations

### Competing interests

The authors declare no conflict of interest.

## Notes

### Competing Interest Statement

The authors have declared no competing interest.

### Summary of Updates

This version of manuscript has been revised to update manuscripts acknowledgement and supplementary file

