## Supplemental File for "Selective degradation of oncogenic extrachromosomal DNA by type I-E CRISPR"

**Supplementary Figures**

**Suppl Figure 1**

COLO320DM

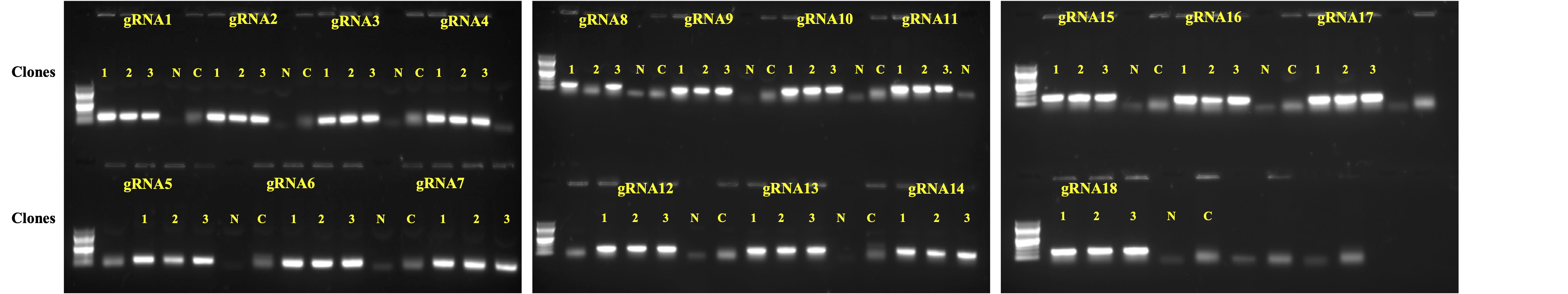

GBM39

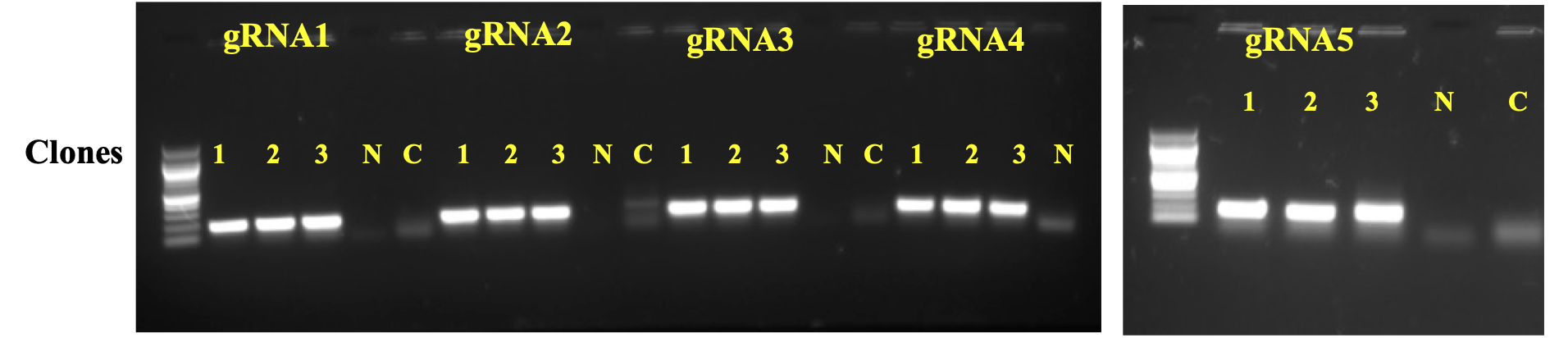

**Suppl. Figure 1-** Agarose gel electrophoresis of colony PCR products to confirm cloning guide RNA oligos into the crRNA plasmid using plasmid backbone primer (M13F) and guide RNA oligo forward primers. C; empty crRNA plasmid control, N; Non-template control. Up; COLO320DM guide RNAs, down; GBM39 guide RNAs.

**Suppl Figure 2**

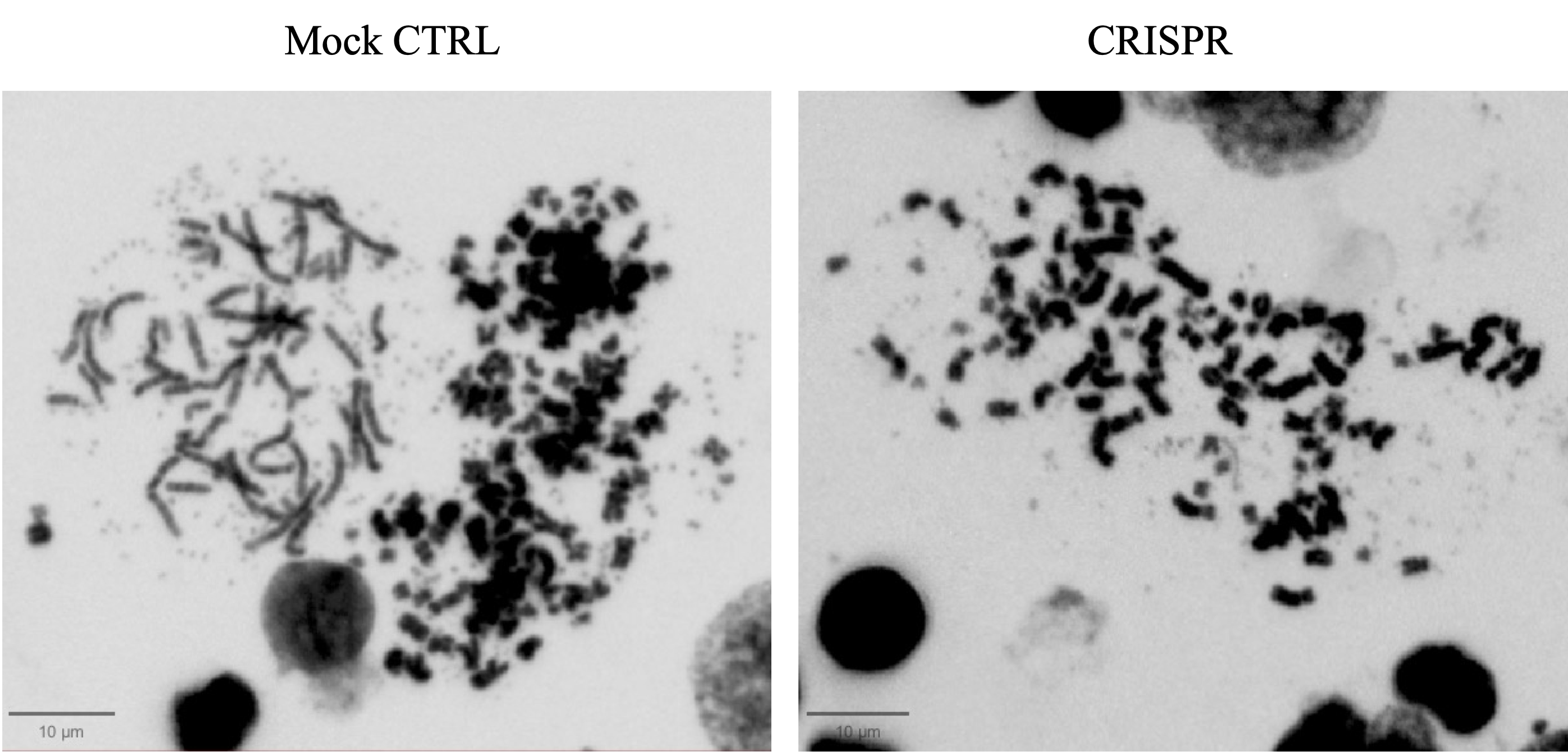

**Suppl. Figure 2.** Metaphase spread of COLO320DM cells 96 hours after treating with type I-E CRISPR system. Cells were stained with DAPI and imaged with Olympus V200 microscope.

**Suppl Figure 3**

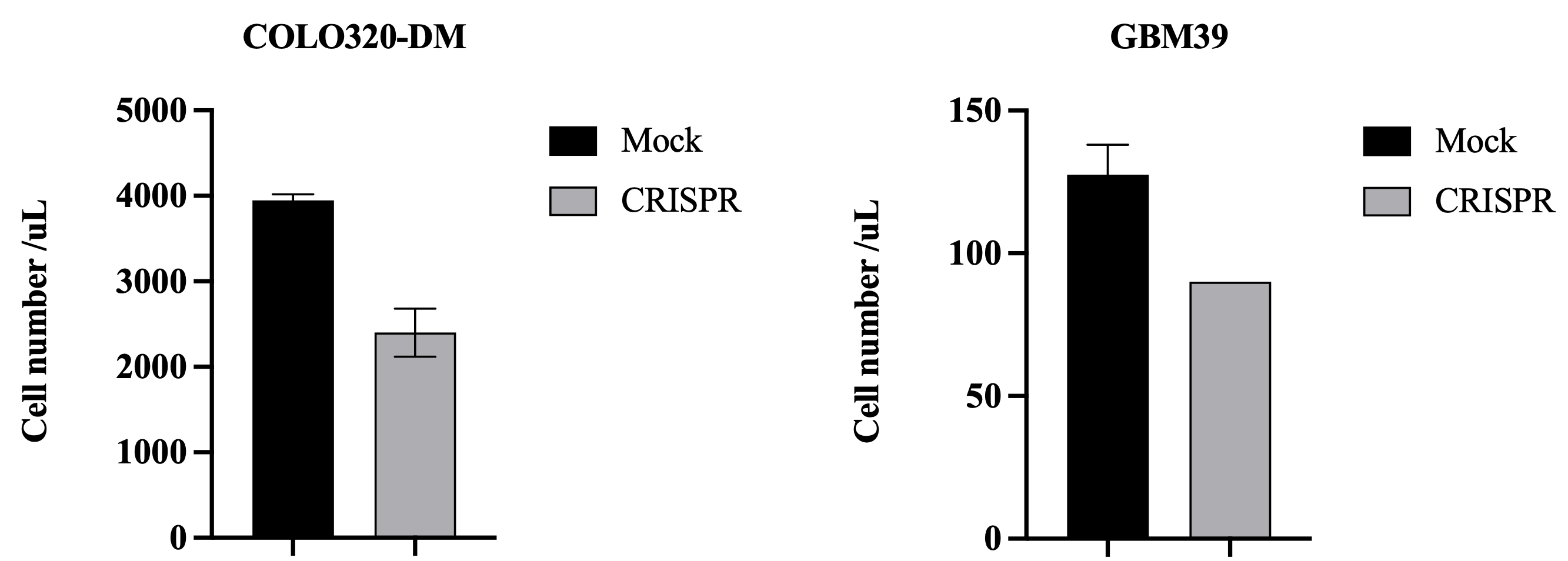

**Suppl. Figure 3.** Cell proliferation of COLO320DM and GBM39 cells 5 days after treatment with type I-E CRISPR system.

**Suppl Figure 4**

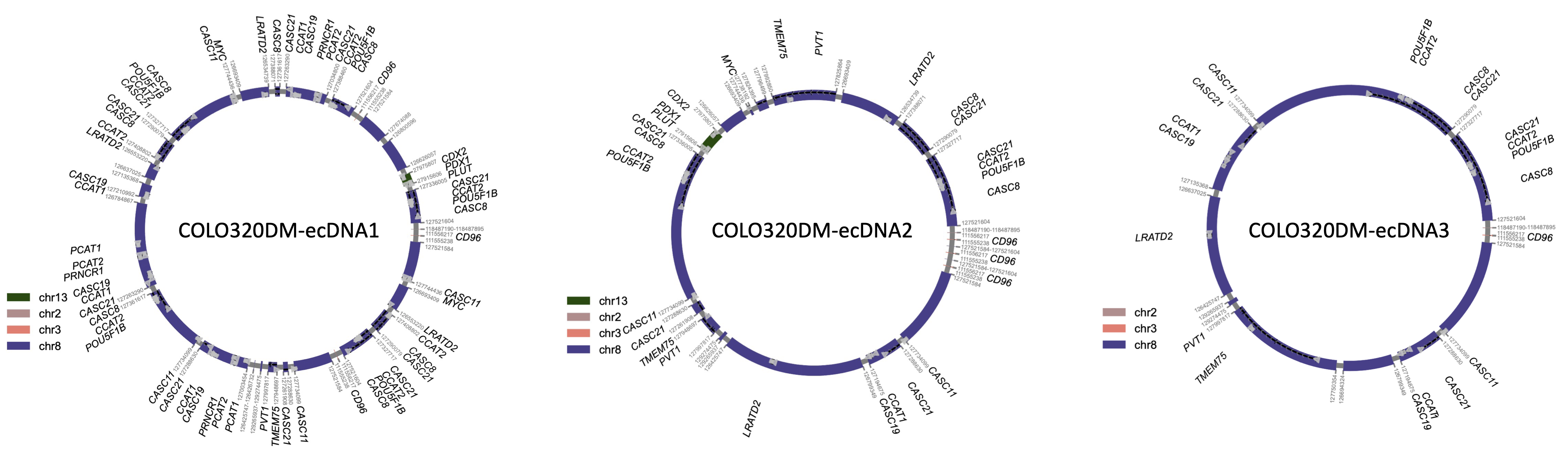

**Suppl Figure 4.** Predicted ecDNA structures for COLO320DM cells

**Suppl Figure 5**

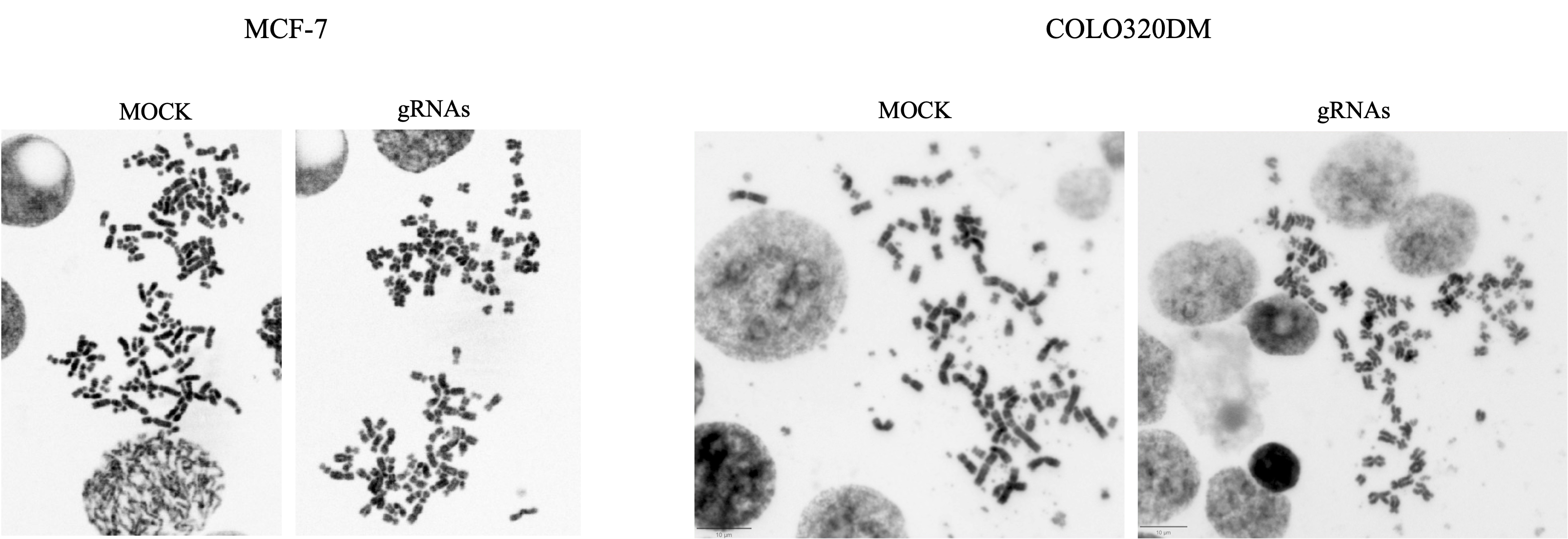

**Suppl Figure 5.** Metaphase spread of MCF7 cells (ecDNA-) and COLO320DM (ecDNA+) 5 days after treating with type I-E CRISPR system. Cells were stained with DAPI and imaged with Olympus V200 microscope.

**Suppl Table 1**

| **Guide RNA** | **PAM** | **Target sequence 5’->3’** |
| --- | --- | --- |
| NeoR-gRNA1 | AAG | AGCATCAGGGGCTCGCGCCAGCCGAACTGTTC |
| NeoR-gRNA2 | ATG | CCGCCGTGTTCCGGCTGTCAGCGCAGGGGCGC |
| COLO320DM_gRNA1 | AAG | CTAGGGTCCACGCGGCCTGGAAGACCCTGATG |
| COLO320DM _gRNA2 | AAG | CAGCAGGAAGGTGTCAAAGGAACTACCATCAG |
| COLO320DM _gRNA3 | AAG | CAGAAAAAGGCAGGGTGGGATCCTGGACCAGG |
| COLO320DM _gRNA4 | ATG | GGTGGATGAATATCATCGCGCCTGGATGTCAA |
| COLO320DM _gRNA5 | ATG | GGTTTGGAACAAATAATGAAGCAAACATTCCA |
| COLO320DM _gRNA6 | ATG | AGATGTGGAGGTTTTTGAGACGGAGTCTCGCT |
| COLO320DM _gRNA7 | AAG | AGACTACCGGTCACTACTACATTTTTTTTTTC |
| COLO320DM _gRNA8 | AAG | TGTAAATTAAATTTCACTCAATGTAGAAGGTG |
| COLO320DM _gRNA9 | ATG | TCAAAAAAGTTCAAGTTTTGGACATAATAAAA |
| COLO320DM _gRNA10 | AAG | AAGAGTCGAATTTTATCCGAGGTGAAGATTAA |
| COLO320DM _gRNA11 | AAG | TGTAAATTAAATTTCACTCAATGTAGAAGGTG |
| COLO320DM _gRNA12 | AAG | AGAACCTTACAAACGAAGTAATAAACAAGGTT |
| COLO320DM _gRNA13 | ATG | GGTTTGGAACAAATAATGAAGCAAACATTCCA |
| COLO320DM _gRNA14 | ATG | CTTTGAAGGACCCAGATATTGCAACTAAGTGG |
| COLO320DM _gRNA15 | ATG | GTAAGCTATATTAAAAAAAAAAGAAGTTGTTA |
| COLO320DM _gRNA16 | ATG | ATCACTGGGCTGGGTAAAGAACTGTGGTACAG |
| COLO320DM _gRNA17 | AAG | CCCTCCATAACTCCTCACCACTGCACTCCAGC |
| COLO320DM _gRNA18 | ATG | GTAAGCTATATTAAAAAAAAAAGAAGTTGTTA |
| COLO320DM _gRNA19 | ATG | AGATGTGGAGGTTTTTTTGAGATGGGGTCTTA |
| COLO320DM _gRNA20 | AAG | GGGTCTTGGTGCGACTTGAGCCCAGGAGTTTG |
| COLO320DM _gRNA21 | ATG | GGATTAGATGTAGACACAGGGAGGAGGTGGCC |
| GBM39_gRNA1 | ATG | GGAAATGTTTCAAAAGTGAGAACTCACTTTGG |
| GBM39_gRNA2 | ATG | GATTTTAAAGCTCTGTTAACTCTCCTTCCTAA |
| GBM39_gRNA3 | ATG | TATGTGTGGTGTGTGTGGTGTGGTGTGTGTAT |
| GBM39_gRNA4 | AAG | AGAATGCAAGTAGTCTACCTCATTTGTAAGAA |
| GBM39_gRNA5 | AAG | AATGTTTACTCCATCTGATGAACGTAAGAGAA |

**Supplementary methods**

**Oligo annealing**

To prepare the guide RNA oligos, a mixture was created by 1 µL of forward oligo, 1 µL of reverse oligo, 1 µL of PNK, 2 µL of 10x T4 DNA ligase, and 15 µL of ddH2O. This mixture was incubated at 37°C for 30 min followed by an incubation at 65°C for 20 min in a thermocycler. For the annealing process of the oligos, a specific thermocycling program was employed: an initial temperature of 95°C for 1 min, followed by a gradual ramp down of 1°C per cycle over the course of 80 cycles, concluding with a final hold at 15°C for 1 min. After annealing, 1 µL of each annealed oligo was utilized for ligation with the guide RNA vector (empty crRNA) using T4 DNA ligase (NEB).

**Guide RNA Cloning**

The pBS-U6-crRNA-empty plasmid was digested with BbsI (NEB) according to the manufacturer's instructions. Annealed guide RNA oligos were then ligated into the plasmid using T4 DNA ligase (NEB) at room temperature for 1 hour and transformed into E. coli DH5-alpha. Positive clones were identified the following day using colony PCR (suppl Fig.1).

**Metaphase chromosome spread**

Coverslips were placed in 12-well plate and soaked in 80% ethanol for 10 min, washed with PBS, coated with poly-D-Lysin and incubated in 37ºC for one h. Then coverslips were washed three times with PBS and air-dried in the tissue culture hood. Cells were added on coverslips and centrifuged for 5 min. Then cells were arrested in metaphase using 100 ng/mL KaryoMAX (Gibco) for 3 h (COLO320DM) or overnight (GBM39). After washing with PBS cells were treated with KCl (75 mM) for 20 min. Samples were then fixed by Carnoy’s fixative (3:1 methanol:glacial acetic acid, v/v).
